# Advances in 16S Taxonomic Classification: Impacts on Species-Level Resolution of Female Bladder Microbiota

**DOI:** 10.64898/2026.10.16.759675

**Authors:** Lisa Karstens, Benjamin Lamp, Juliette Kudrick, Nazema Siddiqui

## Abstract

Bacteria are present in the human urinary bladder, even in the absence of infection. These bacteria comprise the urinary microbiome, or urobiome, and have the potential to promote bladder health or contribute to pathology. Despite its importance, studying the urobiome remains challenging given the high proportion of host to bacterial cells and low microbial biomass. The leading method for studying the urobiome has been 16S rRNA amplicon sequencing. This culture-independent approach provides a cost-effective method for identifying bacteria, but taxonomic classification accuracy depends on three key factors: 1) the variable region targeted; 2) the reference database; and 3) the classification algorithm. In 2021, Hoffman et al. demonstrated that species-level classification of bladder microbiota was possible and that the database used was the most important factor, followed by the variable region. Here, we provide an updated analysis evaluating classification schemes using updated taxonomic databases, most notably GreenGenes2, and expand the analysis to full length 16S gene sequencing. We show that updated taxonomic reference databases improve the ability to identify bladder microbiota at the species level and that long-read sequencing of the full length 16S rRNA gene further improves species identification. These innovations may allow for enhanced discoveries when studying bladder bacteria in the context of bladder disorders.

## Introduction

Microbial communities in the bladder play an important role in urinary tract health, with implications in conditions like urge urinary incontinence^1,2^, or urinary tract infections^3^. Studying these communities remains challenging because the urobiome is notably difficult to culture using traditional methods^4^. 16S rRNA amplicon sequencing has become a key tool for profiling microbial diversity in the bladder and lower urinary tract, offering a cost-effective way to identify bacteria. With this method, bacteria are identified using a taxonomic classification scheme to determine which bacteria an experimental sequence likely belongs to. A taxonomic classification scheme depends on three components:

1. Targeted variable region(s) of the 16S rRNA gene, which act as ‘barcodes’ for differentiating bacteria
2. Classification algorithm, which matches sequences to reference data
3. The reference database, which provides the taxonomy of detected sequences for bacterial identification.

Hoffman et al. (2021) evaluated these components and demonstrated that species-level identification of bladder microbiota can be identified with specific classification schemes^5^. This evaluation demonstrated that the database had that greatest influence on the ability to detect bladder microbiota at the species level, followed by the variable region of the 16S rRNA gene. The two classification algorithms evaluated, Naïve Bayes (NB) and Bayesian lowest common ancestor (BLCA), performed similarly. The comparison of databases (SILVA 132, Greengenes, NCBI 16S RefSeq 2019) revealed the NCBI 16S RefSeq database as the most accurate, outperforming Greengenes, which lacked a wide range of species annotations. While several classifiers showed comparable performance, the database choice dominated accuracy, followed by the choice of amplicon with longer amplicons spanning multiple variable regions (e.g., V1-V3) providing higher resolution over single regions (e.g., V4).

Since Hoffman et al., updates to reference databases have addressed limitations found in the original study. The original Greengenes database, last updated in 2013, underperformed in all metrics compared to its counterparts. Its poor performance in prior analyses, due to lack of coverage of urologically relevant bacteria, has been mitigated by the release of Greengenes2, which is a reconstruction rather than an update. Built from a dataset spanning 15,953 high-quality bacterial genomes, 18,356 full-length 16S sequences, and over 4 million partial sequences (including 2.3 million V4 amplicons), Greengenes2 underwent quality filtering to produce a curated resource of over 21 million sequences with associated taxonomy^6^. SILVA, a leading resource for rRNA sequences and taxonomy, has been maintained through regular updates. Its non-redundant (NR99) dataset (99% sequence identity threshold) ensures high-quality, dereplicated references. While SILVA 132 NR99 contained ∼700,000 sequences, version 138.2 reduced to 510,000 prioritizing accuracy over volume, using improved quality control and dereplication algorithms. The NCBI 16S RefSeq database is smaller, (27,319 sequences, 26,170 bacterial), but remains a great choice for species-level identification due to its stringent curation^7^.

In addition to updates to reference databases, technological advancements have enabled higher accuracy long read sequencing, which has enabled full length 16S rRNA gene sequencing in microbiome research^8^. With these two advancements in mind, we sought to update the Hoffman et al. analysis to reflect the advancements in microbiome science, particularly for the urobiome field which has had substantial growth in the past 5 years.

## Results

Taxonomic classification fidelity, measured by F-measure, varied significantly across reference databases, variable regions, and classification algorithms. Updated databases consistently outperformed their legacy counterparts, with sequence length (single vs. spanning) being a major factor, which was also seen as a major factor in accuracy as seen in Hoffman 2021(**Figure 1**).

**Figure 1:**
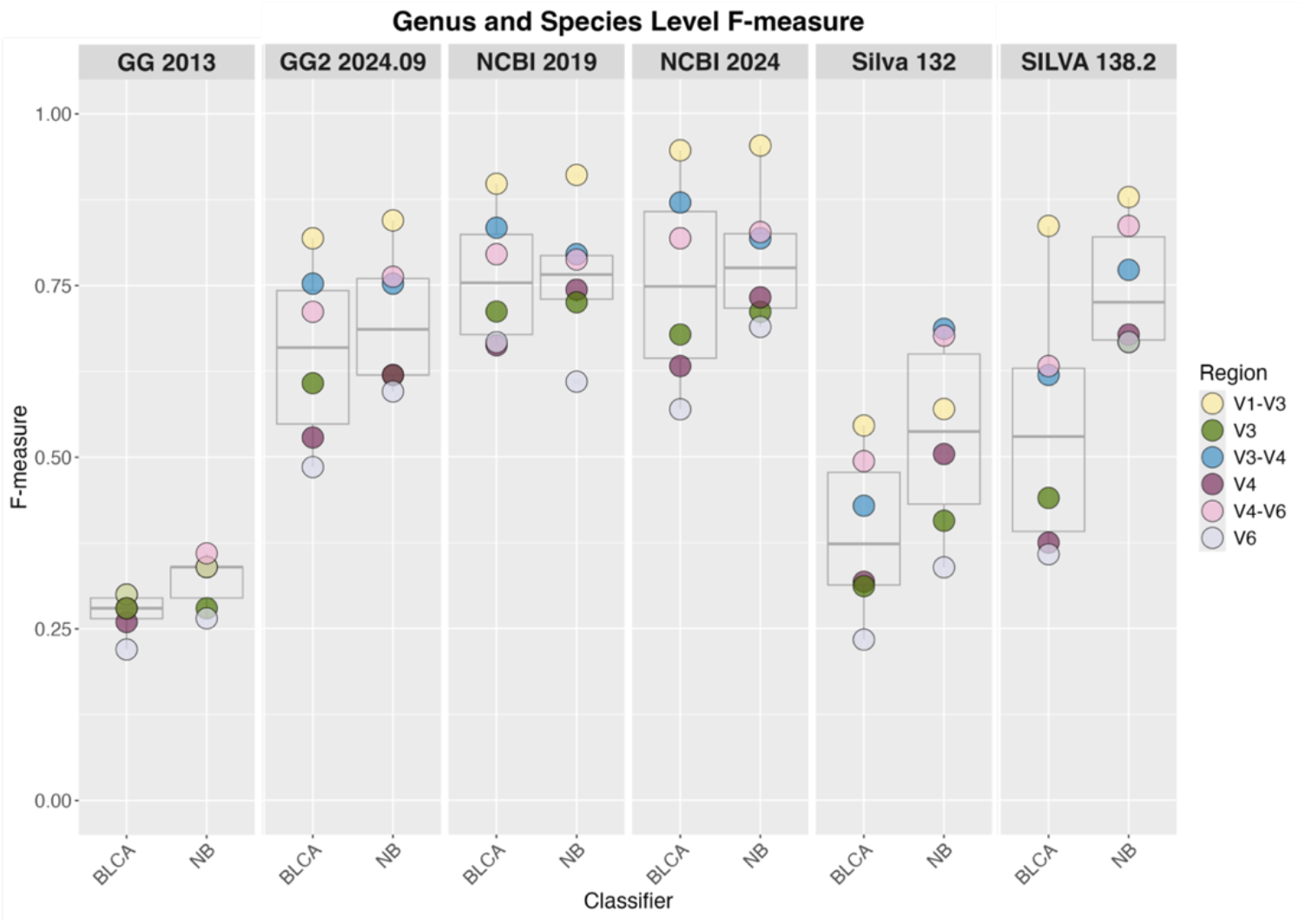
Boxplots demonstrating the spread of F-measures for each classification scheme, faceted by database. Individual dots represent each variable region. Note that shorter length reads of single variable regions are outperformed by multi-region spanning counterparts.

### Database Improvements Over Time

The redeveloped Greengenes2 (GG2) database demonstrated the most improvement. For spanning regions, GG2 2024.09 achieved a mean F-measure of 0.650 with the BLCA classifier, a 122% increase over Greengenes 2013 (F-measure: 0.293). Single-region classifications improved as well (F-measure: 0.540 vs. 0.253 for BLCA)). SILVA’s updates also enhanced performance: SILVA 138.2 achieved a mean F-measure of 0.543 for spanning regions with BLCA, a 40% improvement over SILVA 132 (0.388). The NCBI database maintained high accuracy across updates, with its 2024 version reaching near-perfect classification for full-length sequences (F-measure: 0.967 for BLCA), though single-region performance slightly declined compared to NCBI 2019 (0.626 vs. 0.680).

### Variable Regions and Classifier Performance

Similar to Hoffman et al., variable regions that spanned multiple variable regions (e.g. V4 – V6) performed better than those covering a single variable region (e.g. V4)(F-measure 0.696 – 0.878 compared to 0.391 – 0.711, respectively **Table 1**) and were able to identify more urobiome relevant species (**Figure 2**). We additionally added full-length 16S to our evaluation, which performed better than individual or spanning variable regions in all evaluations (F -measure range 0.861 – 0.967). Additionally, NB and BLCA performed similarly, but had variations based on type of region and database used. For example, for spanning regions, NB outperformed BLCA with Silva (mean F-measure 0.829 vs 0.696, respectively) and Greengenes2 (0.786 vs 0.760), but not NCBI (0.878 vs. 0.866). For full-length data, BLCA slightly outperformed NB with all databases.

**Table 1.** F-measures for Classification Schemes with updated Databases.

| Classifier | Database | Full length | Spanning (mean) | Single (mean) |
| --- | --- | --- | --- | --- |
| **BLCA** | GG2 2024.09 | 0.894 | 0.760 | 0.540 |
| **BLCA** | NCBI 2024 | 0.967 | 0.878 | 0.626 |
| **BLCA** | SILVA 138.2 | 0.939 | 0.696 | 0.391 |
| **NB** | GG2 2024.09 | 0.861 | 0.786 | 0.611 |
| **NB** | NCBI 2024 | 0.96 | 0.866 | 0.711 |
| **NB** | SILVA 138.2 | 0.917 | 0.829 | 0.671 |

**Figure 2.**
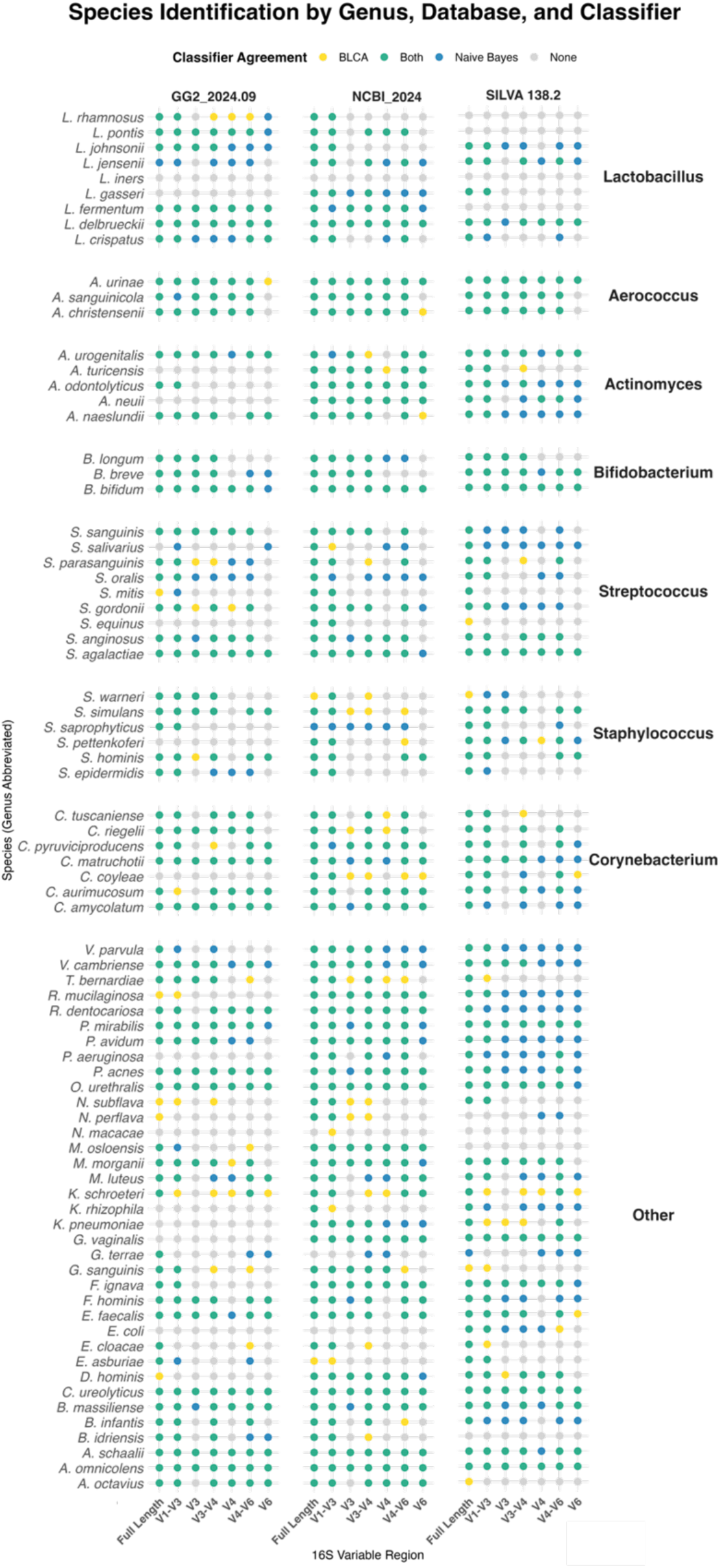
Species identification matrix across 16S rRNA variable regions. The matrix includes 78 bladder bacterial species, grouped by genus, evaluated using different 16S rRNA regions. Each dot represents successful species-level identification for a given region. Full-length 16S performed the best across all classifiers, providing the highest resolution for species-level classification.

### Legacy vs. Current Databases

Greengenes 2013 and SILVA 132 had the lowest F-measures overall (spanning regions: 0.293 and 0.489, respectively), where modern counterparts like GG2 and SILVA 138.2 reduced misclassification errors by up to 171%. The NCBI database proved an exception, maintaining high accuracy across updates (spanning regions: 0.831 in 2019 vs. 0.866 in 2024), though its smaller size limited improvements compared to GG2’s expansive redevelopment.

### Updates to Taxonomy

When comparing new and old datasets or reference materials, it is important to recognize that while sequence data may remain unchanged, nomenclature can evolve. A key example is the reclassification of the *Lactobacillus* genus into 23 novel genera^9^, a change made to better reflect phylogenetic relationships. For clarity in interpretation, we have grouped these genera under *Lactobacillus* for this report. We also observed a misclassification in Greengenes2. When classifying the *Schaalia odontolyticus* sequence, Greengenes2 incorrectly assigns it to the genus *Pauljensenia*, resulting in the label:

Genus: Pauljensenia, Species: Schaalia_odontolyticus

While the full genus_species name in the species assignment is correct, the genus level assignment is inaccurate. This discrepancy stems from the influence of the GTDB (Genome Taxonomy Database) of the Greengenes2 dataset. This is the only instance we encountered, but further discrepancies could remain in the full database.

## Conclusions

We performed an updated benchmarking analysis of Hoffman et al, using updated classification schemes, evaluating the impact of database, classifier algorithms, and variable regions on taxonomic assignment accuracy for bacterial species found in the urobiome. We demonstrate that overall, taxonomy classification of urobiome species is improved with the use of updated databases. We additionally demonstrate that full-length 16S sequencing improves species-level classification. We also noted several updates to taxonomy nomenclature as well as errors in database listings. Both of these issues in taxonomy assignment are important to keep in mind when performing meta-analyses and when comparing results to prior literature. Taxonomic resources such as the *List of Prokaryotic Names with Standing in Nomenclature* (LPSN) or the NCBI Taxonomy Browser, can help with resolving historical discrepancies.

## Methods

### Data Acquisition, Multiple Sequence Alignment and Amplicon Extraction

Bladder-relevant type strain sequences from Thomas-White et al^10^. was obtained from the SILVA v138 database on August 1, 2024. All sequences were compiled into a multi-record FASTA file for processing. Multiple sequence alignment of the compiled FASTA file was performed using T-Coffee (v. default parameters). 16S variable regions were extracted with extract_16s_vr.py script, which relies on primer-annealing coordinates identified in *Escherichia coli* type strain (GenBank accession: EU014689). Primer binding sites were mapped using UGENE, the coordinates were transposed onto the aligned sequences in T-Coffee to extract each target amplicon region.

### Taxonomic Assignment

Naïve Bayes (NB)^11^: Classifications were generated using the assignTaxonomy function of DADA2^12^ with an 80% bootstrap confidence threshold.

Bayesian lowest common ancestor (BLCA)^13^: The BLCA classifier was run using default settings, pointing to the selected reference database.

### Database Construction

To accommodate different classification approaches, two types of reference databases were constructed from the curated type-strain FASTA:

NB Databases: A taxonomy file and a multi-level FASTA file were created for input to Naïve Bayes classification tools. This involved assigning each sequence a hierarchical taxonomic path (e.g., Kingdom;Phylum;Class;Order; Family;Genus;Species).

BLCA Databases: A BLAST-based database was generated using the makeblastdb command (BLAST+ suite).

### Synonym Resolution

Bacterial species synonyms were gathered from the Prokaryotic Nomenclature Up-to-Date (PNU) database on January 5, 2020 and updated July 10, 2024^14^. Taxa assigned were cross-referenced against entries and if the reported name matched either a valid species name or a recognized synonym, the assignment was considered correct.

### Evaluation Metrics

Classifier performance was quantified at the genus and species levels using standard precision, recall, and F-measure metrics:

A true positive (TP) was defined as a correctly classified sequence. False positives (FP) were sequences misclassified. False negatives (FN) were sequences that were either unclassified or assigned below the 80% confidence threshold. When applying synonym corrections, an assignment was considered correct if it aligned with any recognized name or synonym.

## Acknowledgements

This work was supported by the National Institute of Diabetes and Digestive and Kidney Diseases through R01DK138121 and the National Library of Medicine through award R25LM014207 (BL). These awards supported the development of the study and completion of the work presented in this manuscript and do not reflect the views of the funding agencies.

